# Phylogeographic analysis reveals an ancient East African origin of the human herpes simplexvirus 2 dispersal out-of-Africa

**DOI:** 10.1101/2022.01.03.474822

**Authors:** Jennifer L. Havens, Sébastien Calvignac-Spencer, Kevin Merkel, Sonia Burrel, David Boutolleau, Joel O. Wertheim

## Abstract

Human herpes simplex virus 2 (HSV-2) is a globally ubiquitous, slow evolving DNA virus. HSV-2 genomic diversity can be divided into two main groups: an African lineage and worldwide lineage. Competing hypotheses have been put forth to explain the history of HSV-2. HSV-2 may have originated in Africa and then followed the first wave of human migration out of Africa between 50-100 kya. Alternatively, HSV-2 may have migrated out of Africa via the trans-Atlantic slave trade within the last 150-500 years. The lack of HSV-2 genomes from West and Central Africa, combined with a lack of molecular clock signal in HSV-2 has precluded robust testing of these competing hypotheses. Here, we expand the geographic sampling of HSV-2 genomes in order to resolve the geography and timing of divergence events within HSV-2. We analyze 65 newly sequenced HSV-2 genomes collected from primarily West and Central Africa along with 330 previously published genomes sampled over a 47-year period. Evolutionary simulations confirm that the molecular clock in HSV-2 is too slow to be detected using available data. However, phylogeographic analysis indicates that all biologically plausible evolutionary rates would place the ancestor of the worldwide lineage in East Africa, arguing against the trans-Atlantic slave trade as the source of worldwide diversity. The best supported evolutionary rates between 4.2×10^−8^ and 5.6×10^−8^ substitutions/site/year suggest a most recent common ancestor for HSV-2 around 90-120 kya and initial dispersal around 21.9-29.3 kya. These dates suggest HSV-2 left Africa during subsequent waves of human migration out of East Africa.

## Introduction

Human herpes simplex virus 1 and 2 (HSV-1, HSV-2) are globally ubiquitous human pathogens. Both HSV-1 and HSV-2 are slow evolving double-stranded DNA viruses that can infect via oral and genital routes, though HSV-1 is predominantly associated with oral herpes and HSV-2 with genital herpes (1). The consequences of genital herpes infection include increased risk of HIV acquisition and mortality in neonates (2, 3). In 2016, worldwide the prevalence of HSV-2 was 13%; the highest prevalence is in Africa with around 44% of female population and 25% of male population living with HSV-2 (4). HSV-2 has ancient zoonotic origins, whereby proto-humans in Africa were infected by an ape simplexvirus (5–7).

Evolutionarily, HSV-2 is comprised of two major phylogenetic lineages: the African lineage and the worldwide lineage, separated by a long internal branch (8). This relationship suggests HSV-2 diversified in the human population in Africa before migrating out of Africa and dispersing worldwide. It has been hypothesized that HSV-2 followed modern humans out of Africa (8) sometime between 50 and 100 kya (9, 10). Alternatively, it has been suggested that HSV-2 left Africa far more recently, sometime between 1671 and 1792, via the trans-Atlantic slave trade (11) which was active between 1501–1867 CE (12). Phylogeographic analysis of other viruses suggests that the slave trade may have been a conduit for the introduction of many viruses into the Americas (13).

The inability to distinguish between these two competing hypotheses operating on vastly different time-scales arises due to a lack of strong temporal signal in the HSV-2 molecular clock when calibrated using viral sampling dates [i.e. tip dating calibration (14)]. For HSV-2, there is no detectable relationship between the year of viral sampling and the distance from the root of the phylogeny (11, 15). The hypothesis of human migration out-of-Africa as a mechanism for HSV-2 dispersal arises from molecular clock estimates calibrated based on inference of codivergence events deeper in the simplexvirus tree that suggest a genome-wide clock rate on the order of 10^−8^ substitutions/site/year (5, 7, 8, 16). In contrast, the hypothesis that the trans-Atlantic slave trade was the route for HSV-2 worldwide dispersal is based upon a power-law rate decay of the molecular clock, whereby more recent branches have a faster clock rate than older branches (17), with an intercept of 3.3×10^− 3^ substitutions/site/year (11). The absence of a molecular clock calibrated exclusively within HSV-2 leaves these competing hypotheses unresolved.

Importantly, each of these origin theories are associated with different geographic sources within Africa. If HSV-2 accompanied the earliest human migration out-of-Africa, then the MRCA of the worldwide diversity would emanate from East, Central, or Southern Africa, where the first modern human migration began (10). However, if HSV-2 left Africa by the trans-Atlantic slave trade, then this lineage would originate from West or Central Africa—where most Africans enslaved originated. Currently, most available African HSV-2 genomes are from East Africa (8, 11, 18), limiting our ability to formally test these hypotheses.

Here, we clarify the location of the emergence of HSV-2 out of Africa by sequencing 65 HSV-2 genomes from 18 previously unsampled countries, including a majority of viral samples from West and Central Africa. We further narrow down the timing of the dispersal out-of-Africa through simulation and empirical molecular clock analysis to determine which origin scenarios are compatible with a lack of clock-like signal in HSV-2. We show that the worldwide lineage of HSV-2 followed humans from East Africa, but likely postdated the initial out-of-Africa human migration event.

## Results

We sequenced the enriched DNA of 65 new HSV-2 isolates, including 47 with a plausible link to 13 African countries, permitting us to determine sequence across a mean of 77% of the HSV-2 genome (range: 45-91%).

### Maximum likelihood phylogenetic inference

These 65 newly sequenced African HSV-2 genomes were combined with previously published HSV-2 genomes resulting in an alignment of 395 taxa, sampled over 47 years from 37 countries (SI Dataset 1; 1A). The total alignment length was 105,520 nt and recombinant regions were removed.

We inferred a maximum likelihood phylogeny using IQTree2 (19), which supported the existence of two distinct HSV-2 lineages: an African lineage and a worldwide lineage (Fig. 1B). These lineages are separated by a relatively long internal branch with length of 0.009 substitutions/site; the clades are supported with an aBayes of 1.0 (SI Dataset 2). Both midpoint rooting and outgroup rooting using the chimpanzee herpesvirus (ChHV) supports rooting the HSV-2 tree on this long branch, consistent with previous studies (8, 11). The African lineage comprised sequences from West Africa (n=9) and Central Africa (n=4).

**Fig. 1.**
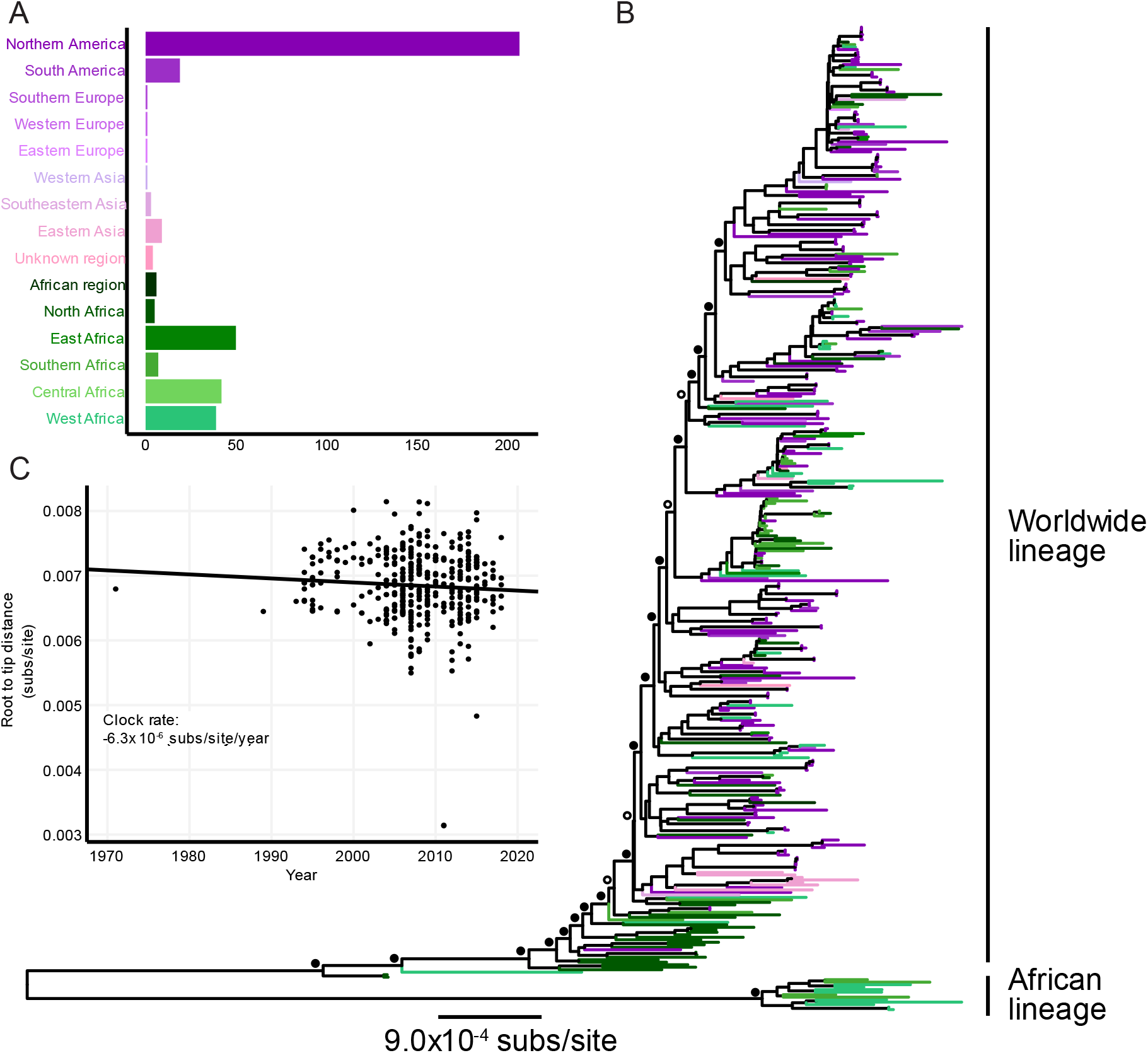
Maximum likelihood (ML) phylogenetic analysis of empirical HSV-2 partial genomes. (A) HSV-2 dataset of 395 sequences by number of samples from each region. (B) ML phylogenetic tree inferred under a GTR+F+R4 model. Tip color indicates geographic region of sampling. The tree is midpoint rooted, though the identical rooting orientation is recovered using the closely related chimpanzee simplexvirus (ChHV). Filled circles indicate nodes with aBayes support >0.95; open circles indicate nodes with aBayes support <0.95. (C) Root-to-tip distance in ML tree versus sampling year with an inferred clock rate of -6.3×10^−6^ substitutions/site/year.

The worldwide lineage contains 382 sequences from all sampled geographic regions, including from West and Central Africa. The basal sequences of the worldwide lineage are primarily East African. The basal structure of the worldwide lineage has strong phylogenetic support (aBayes 0.95-1.00 for the first 8 nodes; Fig. 1). Relative to the basal East African sequences, the crown of this lineage representing global tip diversity has little observable geographic structure (Fig. 1B), likely due to the oversampling of genomes from North America (11, 20, 21).

### Clock rate inference of empirical sequences

In an attempt to estimate the timing of divergence events within HSV-2, we performed molecular clock analysis using TreeTime (22). However, even with the inclusion of HSV-2 genomes sampled over a 47-year timespan, we inferred a negative clock rate (Fig. 1C); the coefficient of determination (R^2^) of the regression between year of sampling and root-to-tip distance was 0.00. These findings indicate a lack of clock-like signal, consistent with previous investigations of the HSV-2 molecular clock using a tip-dating approach (11, 15).

We first explored the possibility that this lack of signal was due either to the phylogenetically recessed basal branches of the worldwide lineage or the long branch separating the worldwide and African lineages. However, analyzing the worldwide lineage after removing these basal taxa did not recover a clock-like signal (Fig. S1).

### Estimation of range for clock rate maximum likelihood

To understand the plausible range of evolutionary rates that would be compatible with the empirical HSV-2 data, we explored the shape of likelihood surface across a series of fixed substitution rates ranging between 10^−8^ and 10^−4^ substitutions/site/year to determine which molecular clock rates have the highest likelihood and what rates are substantially worse. We restricted this analysis to rates faster than 10^−8^ substitutions/site/year, as slower rates would infer an age of HSV-2 older than the age of modern humans. As expected, the slowest rates have the best likelihood scores and the fastest rates have the worst (Fig. 2). Clock rates from 10^−8^ through 3.2×10^−7^ substitutions/site/year are within the 10 log-likelihood points of the maximum likelihood in this analysis. Substitution rates faster than 3.2×10^−7^ substitutions/site/year are a substantially worse fit to the observed data.

**Fig. 2.**
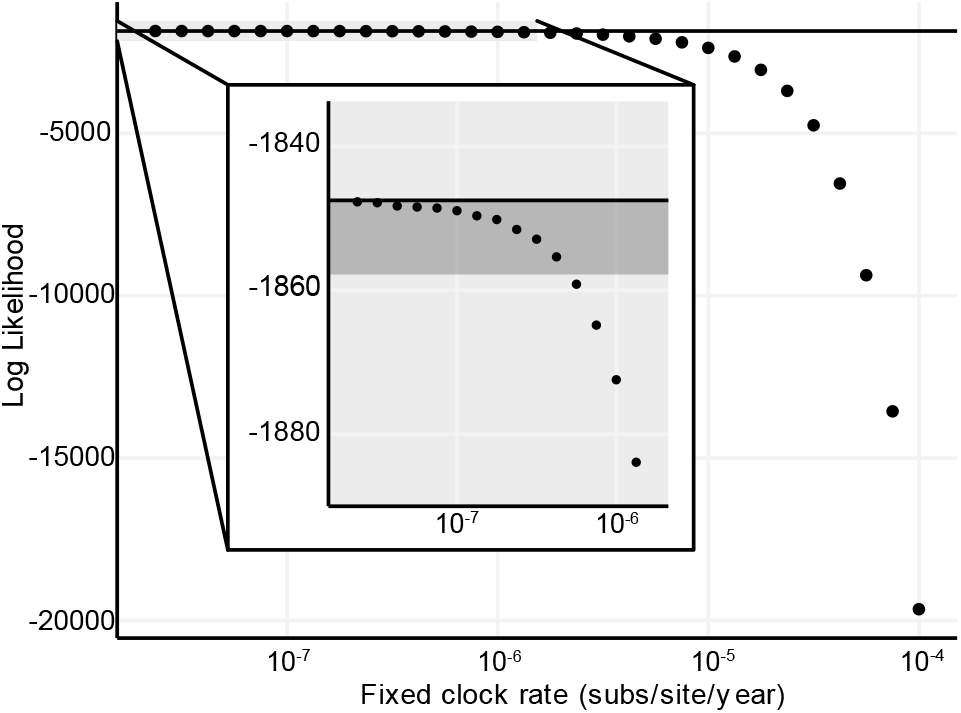
Shape of likelihood surface for clock rate on empirical data. Log-likelihood of HSV-2 time tree with a series of fixed clock rates estimated using the ML topology. The black line is maximum estimated likelihood. The grey shaded box is the area shown in insert. Within the insert, the black line is maximum estimated likelihood at rate 1×10^−8^ substitutions/site/year, and the dark grey box highlights the range 10 points below the maximum likelihood estimate.

We note when we performed the same procedure using a virus with strong molecular clock signal, Ebolavirus from the 2014-2015 West African epidemic (23), the peak of this log-likelihood curve reasonably approximated the ML rate (Fig. S2).

### Estimating clock rate of simulated data

To explore at what substitution rates temporal signal should theoretically be detectable in our HSV-2 dataset, given empirical sampling dates and phylogenetic structure, we simulated HSV-2 genomic data across the inferred ML tree under substitution rates between 10^−8^ and 10^−4^ substitutions/site/year. We then inferred ML trees from these simulated data and re-inferred the clock rate. A simulated rate that had detectable temporal signal would be inconsistent with the empirical data.

We were able to reliably infer clock rates from sequences simulated at 5.6×10^−7^ substitutions/site/year and faster (Fig. 3A), indicating the true underlying HSV-2 substitution rate must be slower than these rates. Based on a breakpoint analysis, for simulated rates slower than 5.6×10^−7^ substitutions/site/year, the mean of the absolute value of inferred clock rates did not capture the simulated rate, unlike faster rates (segmented regression; Fig. S3), suggesting these slower rates are at least consistent with the empirical data.

**Fig. 3.**
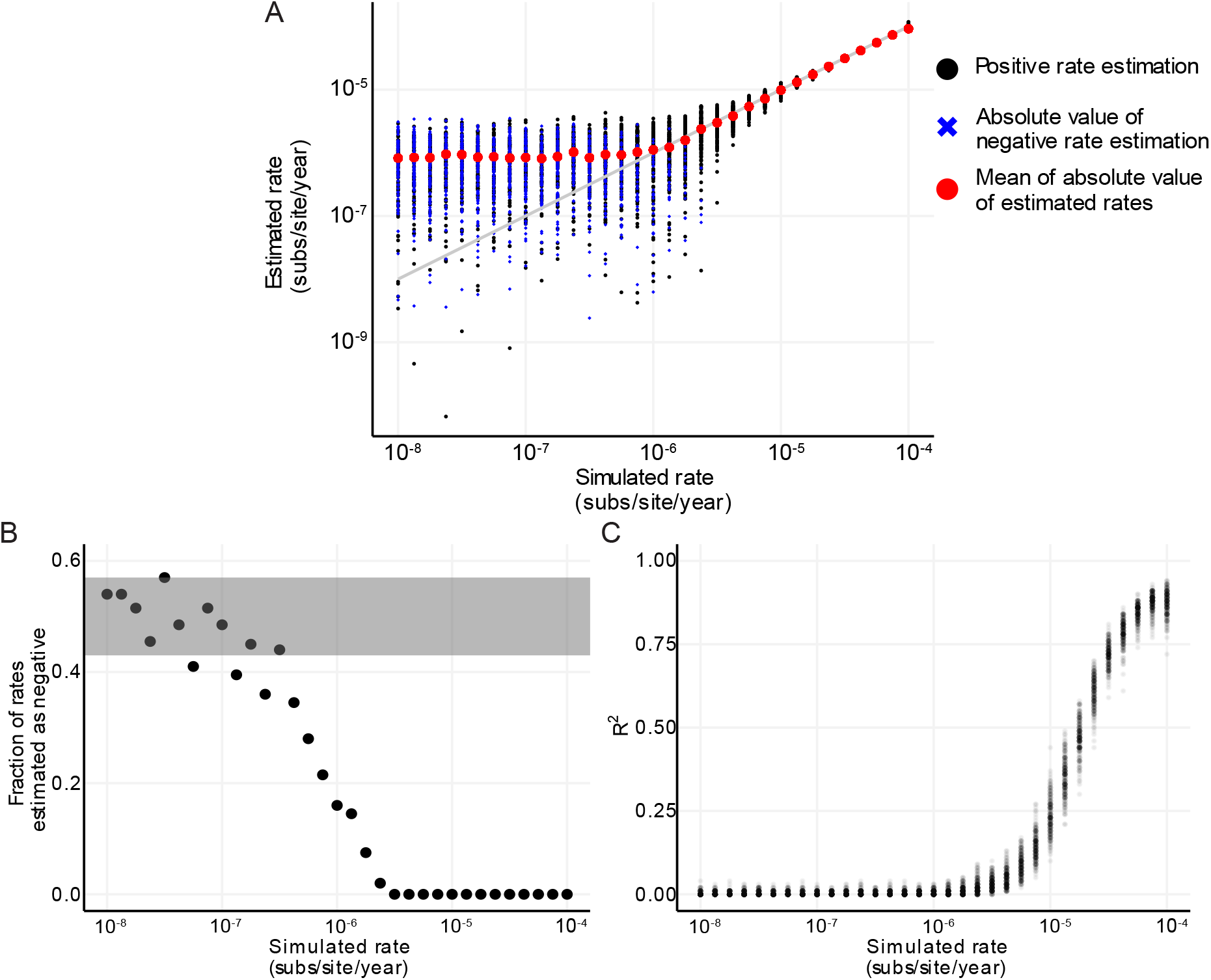
Performance of clock rate inference on simulated data. (A) Inference of clock rate for a fixed input tree with simulated sequences and empirical dates. Black dot represents estimates of positive rates, a blue ‘x’ represents the absolute value of estimates of negative rates, and red dots indicate the mean of absolute value for each fixed rate. Grey line indicates parity of simulated and estimated rate. (B) Negative clock rate analysis of the fraction of total simulations for a fixed rate that have a rate estimated as negative. Grey shading represents range for which a negative estimate is equally probable as positive (binomial test *p*<0.05). (C) R^2^ of root-to-tip regression, a transparent dot represents one replicate, dark dots indicates overlapping estimates for multiple replicates. 200 replicates of simulated were generated under GTR+F+Γ4 model then clock rate estimated on simulated sequences

As in the empirical analysis, the estimated clock rates simulated under slower rates are often negative (Fig. 3B). Thus, sequences simulated under a clock rate which results in an estimated negative rate indicates that the true data could have been produced under that rate. The fastest rate at which more than 5% of trials infer a negative rate on the simulated data is 1.8×10-6 substitutions/site/year (Fig. 3B).

The R2 of the root-to-tip by sample date regression decreases with slower simulated clock rates (Fig. 3C), because as the clock rate, or slope, approaches zero, less of the divergence is explained by sampling date. At rates slower than 2.4 x10-6 substitutions/site/year and 1.3×10-6 substitutions/site/year more than 5% and 50%, respectively, of the R2 were zero (Fig. 3C). The simulations of slower rates are consistent with the root-to-tip regression of the empirical HSV-2 dataset, which had an R2 of zero.

The R^2^ of the root-to-tip by sample date regression decreases with slower simulated clock rates (Fig. 3C), because as the clock rate, or slope, approaches zero, less of the divergence is explained by sampling date. At rates slower than 2.4 ×10^−6^ substitutions/site/year and 1.3×10^−6^ substitutions/site/year more than 5% and 50%, respectively, of the R^2^ were zero (Fig. 3C). The simulations of slower rates are consistent with the root-to-tip regression of the empirical HSV-2 dataset, which had an R^2^ of zero.

### Phylogeographic analysis indicates East African origin of worldwide lineage

We performed phylogeographic reconstruction in TreeTime across a range of substitution rates (between 10^−8^ and 10^−4^ substitutions/site/year) to determine the ancestral location of HSV-2 and out-of-Africa migration source region. This range of substitution rates accounts for scenarios in which the time of most recent common ancestor (TMRCA), or root age, for HSV-2 is older than modern humans, 616 kya at 1.0×10^− 8^ substitutions/site/year, to the similarly unrealistic 84 years ago at 1.0×10^−4^ substitutions/site/year (Table 1). The youngest root age not rejected by any of our clock-rate simulation analyses was 19 kya, assuming 3.2×10^−7^ substitutions/site/year (Fig. 4B). The youngest root age consistent with the most permissive simulation analysis (i.e., >5% R^2^ above 0; at 2.4×10^−6^ substitutions/site/year) was 2.6 kya, or 556 BCE (Fig. S4). Root ages of 109 and 146 kya, that are consistent with a previously identified split in modern humans in Africa around 110-160 kya (10), were obtained assuming a substitution rate of 4.2×10^−8^ and 5.6×10^−8^ substitutions/site/year, respectively (Fig 4A).

**Table 1.** Age of root and out-of-Africa dispersal across clock rates.

| Substitution rate<sup>1</sup> | Root age<sup>2</sup> | Out-of-Africa age<sup>2</sup> | Support for East African source<sup>3</sup> | Molecular clock analyses<sup>4</sup> |
| --- | --- | --- | --- | --- |
| $1.0 \times 10^{-8}$ | 616 kya | 124 kya | 0.976 | Not rejected |
| $1.3 \times 10^{-8}$ | 455 kya | 93 kya | 0.977 | Not rejected |
| $1.8 \times 10^{-8}$ | 341 kya | 70 kya | 0.976 | Not rejected |
| $2.4 \times 10^{-8}$ | 256 kya | 52 kya | 0.977 | Not rejected |
| $3.2 \times 10^{-8}$ | 192 kya | 39 kya | 0.977 | Not rejected |
| $4.2 \times 10^{-8}$ | 146 kya | 29, kya | 0.977 | Not rejected |
| $5.6 \times 10^{-8}$ | 109 kya | 22 kya | 0.977 | Not rejected |
| $7.5 \times 10^{-8}$ | 81 kya | 16 kya | 0.977 | Not rejected |
| $1.0 \times 10^{-7}$ | 61 kya | 12 kya | 0.977 | Not rejected |
| $1.3 \times 10^{-7}$ | 45 kya | 9.3 kya | 0.977 | Not rejected |
| $1.8 \times 10^{-7}$ | 34 kya | 6.9 kya | 0.977 | Not rejected |
| $2.4 \times 10^{-7}$ | 26 kya | 5.2 kya | 0.978 | Not rejected |
| $3.2 \times 10^{-7}$ | 19 kya | 3.9 kya | 0.977 | Not rejected |
| $4.2 \times 10^{-7}$ | 14 kya | 2.9 kya | 0.977 | Rejected by some |
| $5.6 \times 10^{-7}$ | 11 kya | 2.2 kya | 0.976 | Rejected by some |
| $7.5 \times 10^{-7}$ | 8.1 kya | 1.7 kya | 0.977 | Rejected by some |
| $1.0 \times 10^{-6}$ | 6.1 kya | 1.2 kya | 0.976 | Rejected by some |
| $1.3 \times 10^{-6}$ | 4.6 kya | 935 ya | 0.976 | Rejected by some |
| $1.8 \times 10^{-6}$ | 3.4 kya | 702 ya | 0.976 | Rejected by some |
| $2.4 \times 10^{-6}$ | 2.6 kya | 528 ya | 0.976 | Rejected by some |
| $3.2 \times 10^{-6}$ | 1.9 kya | 401 ya | 0.975 | Rejected by all |
| $4.2 \times 10^{-6}$ | 1.4 kya | 303 ya | 0.976 | Rejected by all |
| $5.6 \times 10^{-6}$ | 1.1 kya | 230 ya | 0.977 | Rejected by all |
| $7.5 \times 10^{-6}$ | 826 ya | 175 ya | 0.977 | Rejected by all |
| $1.0 \times 10^{-5}$ | 616 ya | 133 ya | 0.980 | Rejected by all |
| $1.3 \times 10^{-5}$ | 463 ya | 104 ya | 0.982 | Rejected by all |
| $1.8 \times 10^{-5}$ | 353 ya | 80 ya | 0.985 | Rejected by all |
| $2.4 \times 10^{-5}$ | 267 ya | 64 ya | 0.989 | Rejected by all |
| $3.2 \times 10^{-5}$ | 205 ya | 55 ya | 0.990 | Rejected by all |
| $4.2 \times 10^{-5}$ | 159 ya | 52 ya | 0.980 | Rejected by all |
| $5.6 \times 10^{-5}$ | 125 ya | 50 ya | 0.944 | Rejected by all |
| $7.5 \times 10^{-5}$ | 101 ya | 49 ya | 0.827 | Rejected by all |
| $1.0 \times 10^{-4}$ | 84 ya | 49 ya | 0.540 | Rejected by all |

<sup>1</sup> Substitutions/site/year
<sup>2</sup> Prior to most recently sampled virus, 2018
<sup>3</sup> Maximum inferred probability for East African ancestor prior to out-of-Africa dispersal
<sup>4</sup> Empirical and simulation analyses described in Figure 5

**Fig. 4.**
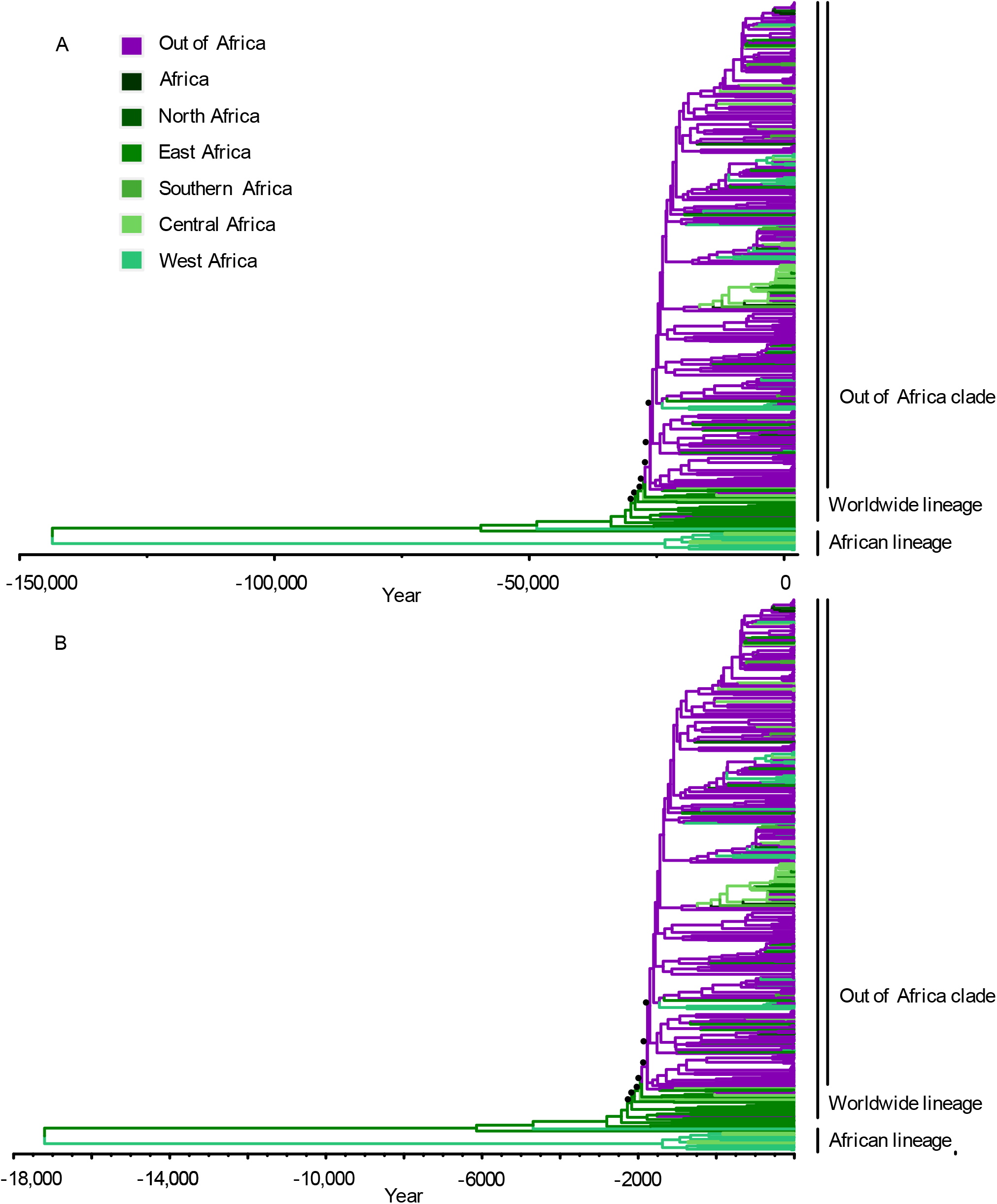
Phylogeography of HSV-2 supports East African origin of out-of-Africa dispersal. (A) Time tree inferred using the evolutionary rate of 4.2×10^−8^ substitutions/site/year. (B) Time tree inferred with the fastest rate supported by all clock analyses: 3.2×10^−7^ substitutions/site/year. ML tree topology scaled to time tree, colored by plurality of support in mugration analysis for each region. Time scale represents the calendar year; negative years are BCE. Black dots indicate nodes used in out-of-Africa weighted age calculation (see Fig. S1).

For every substitution rate explored, the reconstructed ancestral location of the African lineage was in West Africa and the location of the worldwide lineage was in East Africa (Table 1). The support that the ancestral location of the worldwide lineage is East Africa increases with slower substitution rates, exceeding 0.95 for all evolutionary rates that were not rejected by the previous simulation analyses.

The most recent estimate for HSV-2 transitioning from East Africa to out-of-Africa that was based on a rate compatible with all our simulation results is 3.9 kya, or 1883 BCE (Table 1). The out-of-Africa age estimate that is consistent our most permissive rejection strategy was 528 years ago, or 1490 CE; however, the origin of this event was still inferred to be in East Africa. The estimates for out-of-Africa dispersal, assuming a substitution rate of either 4.2×10^−8^ or 5.6×10^−8^ substitutions/site/year which are associated with root age over 100 kya, is 29.3 or 21.9 kya, respectively (Fig. 4). The inference of East African origin for the worldwide dispersal is robust to geographic partitioning schemes of African and non-African locations (Table S1).

## Discussion

Understanding the geographic spread of human pathogens, both ancient and recent, is important because it provides insight into what conditions lead to viruses going extinct or establishing in human populations and improves our understanding and ability to predict the dynamics of future epidemics (24–26). By expanding the geographic breadth of HSV-2 sampling and establishing the sensitivity of molecular clock inference, we conclude that HSV-2 most likely accompanied humans migrating from East Africa between 21.9 and 29.3 kya (Fig. 5), post-dating the original human out-of-Africa migratory event by tens of thousands of years (10). The rates that infer these migration dates also infer date of initial divergence in HSV-2 consistent with previous TMRCA estimates for HSV-2 between 90-120 kya (7, 8). It is therefore unlikely that current worldwide HSV-2 diversity is the result of migrating with modern humans during the original out-of-Africa migration between 50 and 100 kya, as this scenario would place the TMRCA of HSV-2 between 250 and 500 kya, predating modern humans as well as previous HSV-2 age estimates. We acknowledge it is possible that HSV-2 was present during the original out-of-Africa migration and was later replaced by viruses from a subsequent migratory wave.

**Fig. 5.**
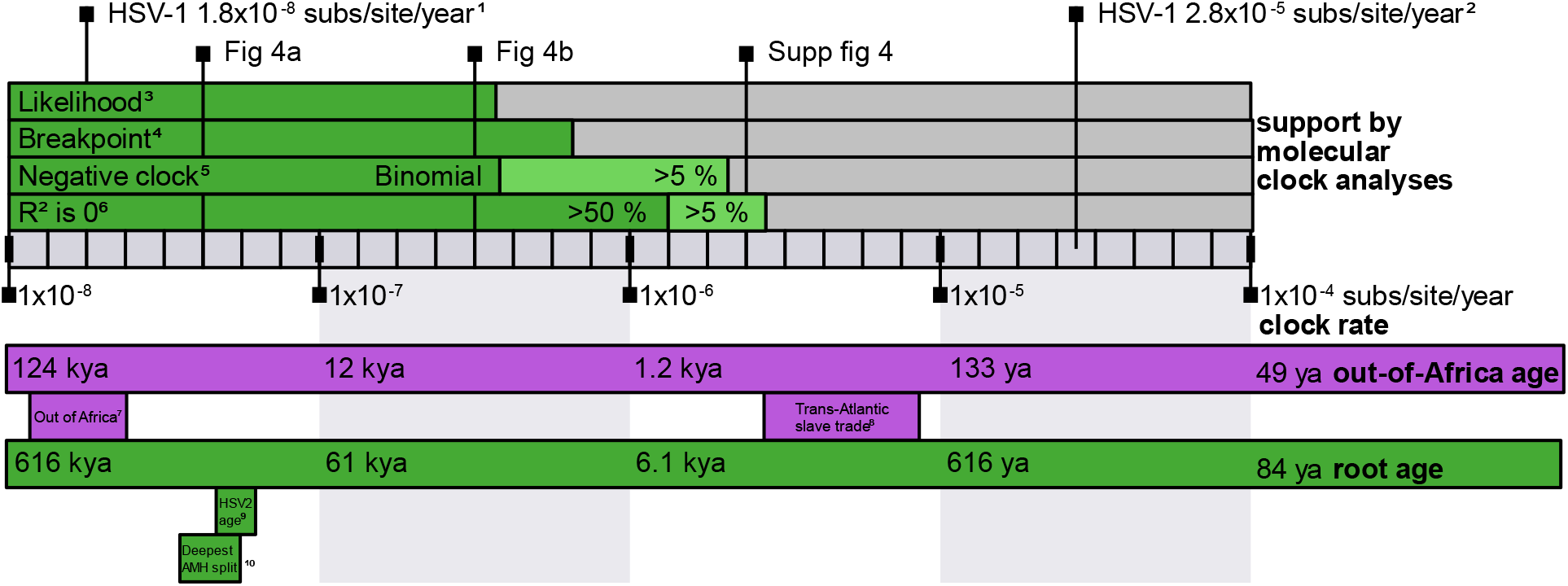
HSV-2 statistical support and dispersal timing across possible molecular clock rates. Upper panel: dark green represents clock rate is supported by specific analysis, light green represents intermediate support, while grey represents the rate is not supported by analysis. Estimates of HSV-1 clock rate indicated at ^(1)^1.8×10^−8^ (16) and ^(2)^2.8×10^−5^ (11) substitutions/site/year. ^(3)^ Likelihood of time tree for empirical data with fixed rates slower than 3.2×10^−7^ substitutions/site/year were within 10 points of maximum log likelihood.^(4)^ Mean of absolute value of clock rates estimated from sequences simulated under rates slower than 5.6×10^−7^ substitutions/site/year are not consistent with true simulated rate.^(5)^ Negative clock rate equally probable to be estimated as positive clock rate (binomial test; *p*<0.05) when simulated under rates slower than 3.2×10^−7^ substitutions/site/year, and more than 5% of clock rate estimates are negative for data simulated under clock slower than 1.8×10^−6^ substitutions/site/year. ^(6)^ R^2^ is zero for >50% and >5% of root to tip regressions for data simulated under that rates slower than 1.3×10^−6^substitutions/site/year and 2.3×10^−6^substitutions/site/year, respectively. (lower panel) Age of nodes across molecular clock rates. Purple boxes indicate ages and events with respect to out-of-Africa dispersal. ^(7)^Timing of first human migrations out of Africa estimated to be 50-100 kya (10). ^(8)^ Timing of trans-Atlantic slave trade ca. 1501–1867 CE (12). Green boxes indicate ages and events with respect to root node and initial HSV-2 divergence. ^(9)^Age of HSV-2 estimated to be 90-120kya, based on internal node calibrations (7, 8). ^(10)^Timing of deepest anatomically modern human split inferred to be 110-160 kya(10).

We conclude that a scenario in which HSV-2 left Africa via the trans-Atlantic slave trade between 1500–1850 CE is inconsistent with both molecular clock and phylogeographic inference. Out-of-Africa migrations events after ca. 1600 CE would necessitate a viral substitution rate that would produce some clock-like signal using tip-date calibrated molecular clock (*≥*3.2×10^−6^ substitutions/site/year). Our clock-signal breakpoint analysis (Fig. S3) indicates that an out-of-Africa event post-dating 191 BCE (i.e., substitution rate 5.6×10^−7^) would necessitate a molecular clock detectable in this dataset. Any internally consistent molecular clock rate places the origin of this out-of-Africa migration in East Africa, the opposite side of the continent from where the majority of Africans enslaved as part of the trans-Atlantic slave trade originated. There is evidence of other viral pathogens being transported to the Americas from Africa during this time (13), we note that it is possible that HSV-2 was transported to the Americas via the trans-Atlantic slave trade after the initial worldwide dispersal. However, we find the timing and geography for the source of HSV-2 worldwide dispersal are incompatible with the slave trade.

A possible explanation for the previous inference of a younger age of the HSV-2 out-of-Africa dispersal is the misapplication of substitution rate correction for purifying selection. As viruses evolve for long periods of time, purifying selection can lead to underestimation of branch lengths due to mutational saturation (27, 28). In RNA viruses, this effect is most prominent on internal branches >0.1 substitutions/site. Various remedies to this problem have been proposed (5, 29), including the rescaling of branch lengths by a power law decay rate, proposed by Aiewsakun and Katzourakis (2016). This latter method was applied to HSV-2 by Forni et al. (11). Importantly, decay of observed rates is not expected until saturation starts and should therefore not apply to trees with relatively short branch lengths, like our HSV-2 phylogeny with a tree height of 0.0083 substitutions/site. We note that a new formulation of the power law decay rate model by Ghafari et al. (30) now describes rate variation through time with a sigmoid curve, acknowledging a more biologically plausible constant evolutionary rate near the tips of viral evolutionary trees with rate decay occurring only deeper in the tree. In other words, although rate decay is almost certainly relevant to deeper divergence events within the simplexvirus phylogeny that show evidence of saturation (5), we believe that HSV-2 is young enough for traditional phylogenetic inference to properly associate branch lengths with the molecular clock.

Although the geographic origins of the worldwide lineage of HSV-2 have now come into focus, the path the virus took upon leaving Africa is still unclear. Modern-day sampling is strongly biased towards North America, and human migration in the modern era likely obscures ancient signal. Previous analysis of HSV-2 geographic structure has primarily partitioned samples into continent based regions (11, 20, 21). Increased sampling across global regions could improve geographic signal of migration pattern of HSV-2, once it migrated out of Africa. The identification of ancient herpes virus sequences (including HSV-1) in a few archeological specimens (31) may foreshadow the possibility of reconstructing ancient HSV-2 genomes, that could be used to disentangle this geography, as well as to inform a tip-dated calibration for this virus.

## Materials and Methods

### HSV-2 data set and sequence alignment

Clinical isolates were recovered from patients diagnosed with genital herpes at La Pitié Salpêtrière – Charles Foix University Hospital (Paris, France) who self-reported overseas origin. HSV-2 isolates were used because of shortages of primary clinical sample material. Isolates were obtained by propagation in subconfluent Vero cell monolayers (32). A limited number of passages was necessary for the generation of viral stocks.

We sequenced 65 HSV-2 samples collected 2012-2018 primarily from West and Central Africa. In brief, we extracted DNA from supernatant using the QIAmp Viral RNA Mini Kit (Qiagen, Hilden, Germany) and measured DNA content with a Qubit (Thermo Fischer Scientific, Waltham, MA, USA). HSV-2 copy numbers were determined using a quantitative PCR (qPCR) assay (8). We then prepared dual indexed-libraries using 1ng DNA extract using the Nextera XT Library Preparation Kit and the Nextera XT Index Kit (Illumina, San Diego, CA, USA), which we pooled so individual libraries would all bring in about the same number of HSV-2 genome copies (according to the qPCR results). The pool was subjected to a double round hybridization capture with a MYbaits Custom Target Enrichment Kit designed to cover the whole genomes of HSV-1 (NC_001806), HSV-2 (NC_001798) and varicella zoster virus (VZV, NC_001348; *Human herpesvirus 3*) (Daicel Arbor Biosciences, Ann Arbor, MI, USA). For both rounds, we followed manufacturer’s instructions with the exception that only one fourth of the recommended bait quantity was used. We sequenced the capture product on MiSeq and NextSeq 550 platforms (Illumina) using the MiSeq Reagent Kit v3 (600 cycles; Illumina) and NextSeq Mid Output Kit v2.5 (300 cycles; Illumina), generating a total of 61,926,628 reads.

Low quality bases and adapter sequences were removed from raw reads using Trimmomatic (33) and overlapping reads were merged using ClipAndMerge (34). We then mapped trimmed reads to the HSV-2 RefSeq genome (NC_001798) from which we had removed the TRL and TRS regions (35). We finally sorted mapping files and removed duplicate reads using the SortSam and MarkDuplicates tools from Picard (http://broadinstitute.github.io/picard/). We generated consensus sequences form the maps using Geneious Prime (36), calling bases at positions covered by at least 20 unique reads and for which more than 95% of the reads were in agreement.

Additionally, we downloaded all available HSV-2 sequences of length greater than 100kbp from GenBank in November 2019. Potentially engineered (n=5) and dubious sequences (n=10) were removed. These HSV-2 sequences along with, ChHV (NC_023677) and HSV-1 (NC_001806) were aligned with MAFFT v7.307 (37). Regions with more than 10% of sequences containing gaps were removed with Geneious. Invariant sites of the alignment were removed, resulting in an alignment of 25,809 sites of ChHV, HSV-1, and HSV-2 sequences. Using variant sites only was done to speed up recombination analysis. Recombinant regions were detected with RDP4 (38), using five recombination detection methods (RDP, GENECONV, MaxChi, BootScan and SiScan) and validating recombination events identified by *≥*2 methods. Recombinant fragments of sequences were removed from the alignment.

### Phylogenetic and initial temporal analysis

Subsequent analysis was done on only HSV-2 sequences with the year of sample collection. Resulting in a final HSV-2 dataset of 395 sequences, with 5,648 variable sites, which are sampled globally across 47 years, from 1971 to 2018 (SI Dataset 1). Constant sites were appended to the variable site alignment, in proportion to the base frequencies of whole genomes. This resulted in an alignment of 105,420 sites.

A maximum likelihood phylogeny for the HSV-2 dataset was made using IQtree2 v2.0.4 (19), with a GTR+F+R4 substitution model selected by model testing using ModelTesting with cAIC on the initial alignment (39). We used TreeTime v0.7.6 (22) to perform molecular clock inference on HSV-2 using tip date calibrations and the input tree with fixed topology, root placement, and branch lengths. Default TreeTime parameters were used, except for fixing the root, using input branch lengths, and clock filter was set to 0 (no filtering). The root was fixed, as this rooting is supported by including ChHV as an outgroup and is consistent with previous studies (8). Input topology and branch lengths were used to because joint optimization is not stable without this approach (40). With the lack of clock like signal it is was not appropriate to filter based on clock variation.

### Likelihood of fixed clock rates with empirical data

A range of fixed substitution rates were applied, and the log likelihood was calculated for the time trees with TreeTime. We used a fixed input tree, empirical sample dates, and no clock filter. The rates used were 1×10^−8^ substitutions/site/year to 1 ×10^−4^ substitutions/site/year, with a resolution of 8 rates distributed evenly across the log space across each order of magnitude. This distribution exceeds the range of previous estimates of clock rates for HSV-1 (11, 16).

To validate the interpretation of the likelihood surface, we performed a similar process with a previously published dataset of 1,610 ebolavirus sequences from the 2014-2015 West African Ebola epidemic (23), which has detectable temporal signal using tip-date calibrations. The Ebola dataset was analyzed with rates from 1×10^−1^ substitutions/site/year to 1 ×10^−5^ substitutions/site/year, with a resolution of 3 rates for each order of magnitude spread evenly across log space (Fig. S2).

### Clock rate estimates of simulated data

In order to understand what clock rate would result in sufficient temporal signal to detect the molecular clock on the empirical HSV-2 dataset (i.e., range of sampling dates, and the length of the sequences, and tree topology), we simulated sequence data across a range of substitution rates, 10^−8^ to 10^−4^ substitutions/site/year with 8 rates across each order of magnitude. We first inferred a time tree with a fixed clock rate based on our topology using TreeTime and then scaled this tree to substitutions/site with the clock rate. Sequences were simulated on the rescaled tree with Seq-Gen v1.3.4 (41). We performed 200 replicates of sequence evolution and subsequent analysis.

Simulation was done with the nucleotide substitution model parameters inferred by IQtree2, under a GTR+F+Γ4 model on the HSV-2 dataset. Branch lengths were estimated on the fixed empirical topology using the simulated sequences with IQtree2 using GTR+F+Γ4 model. Although the empirical tree inference was performed with free rate variation, we used gamma rate variation in the simulations and subsequent tree inference to maintain consistency between simulation, tree and clock inference. The clock rate and R^2^ of the root-tip-regression of the simulated data was then estimated with TreeTime, using previous parameters.

The detection of temporal signal was assessed though different methods using R (42). The rate at which there was a change in the relationship between the mean absolute value of estimated rate and the simulated rate was inferred by a linear regression break point analysis (R segmented v. 1.2-0).

The point at which the estimated rate reflected the empirical data, in that the estimated clock rate was negative, was determined by two cutoffs. The first cutoff method is the fastest rate at which more than 5% of estimated rates on simulated data were negative. The second cutoff is the fastest rate at which a negative estimate is as likely as a positive estimate, determined by binomial test (*p*<0.05).

### Phylogeographic ancestral state reconstruction

We used TreeTime mugration to reconstruct the ancestral geographic state using default parameters, empirical time trees built with different fixed clock rates and the region of sampling for each tip. Samples with country histories were partitioned into regions of North Africa, West Africa, East Africa, Central Africa, Southern Africa, and ‘not Africa’. Six samples had no country of origin, but are known to be sampled from Africa, and were partitioned into a broad ‘Africa’ region for phylogeography analysis. Additionally, 4 samples were from an unknown location and were indicated as ‘missing’ in the mugration analysis.

Robustness analysis of the phylogeographic reconstruction was done by repeating previous mugration analysis with a merged West, Central, and Southern Africa region. Additional robustness analysis split ‘not Africa’ into worldwide subregions, and reconstruction was done with North Africa, West Africa, East Africa, Central Africa, Southern Africa, Africa, Eastern Europe, Southern Europe, Western Europe, Northern America, South America, Southeastern Asia, Western Asia, and unknown.

The points of particular interest in the trees are first the age of the root, which represents the age of HSV-2 diversification, and second the age of the most recent common ancestor (MRCA) of the worldwide diversity within the worldwide lineage, which represents the maximum age for the dispersal of HSV-2 out of Africa, assuming an African origin for the worldwide lineage. On the fixed topology, ancestral geographic reconstruction along the basal backbone of the worldwide lineage had majority of support for the African region that is the source of HSV-2 worldwide diversity which decreases until the majority of support is for the out of Africa region (Fig. S5). We defined the out of Africa age as the average age of the nodes that define basal backbone of the worldwide lineage, weighted by a factor of 1-absoluteValue(support for source African region – support for out of Africa region). This factor results in low weighting to ages that have high support for one region or the other and the highest weight being applied to the age where there is support for both the source and out of African region, which is the most likely time of migration.

Age is reported in number of years ago, relative to 2018, the most resent sample date.

## Supporting information

datasetS1_metadata

datasetS2_treefile

## Acknowledgments

We thank Richard Neher for his guidance on reporting and interpreting likelihoods in TreeTime. This research was supported by National Institute of Health R01 (AI135992).

## Supplementary information

**Fig. S1.**
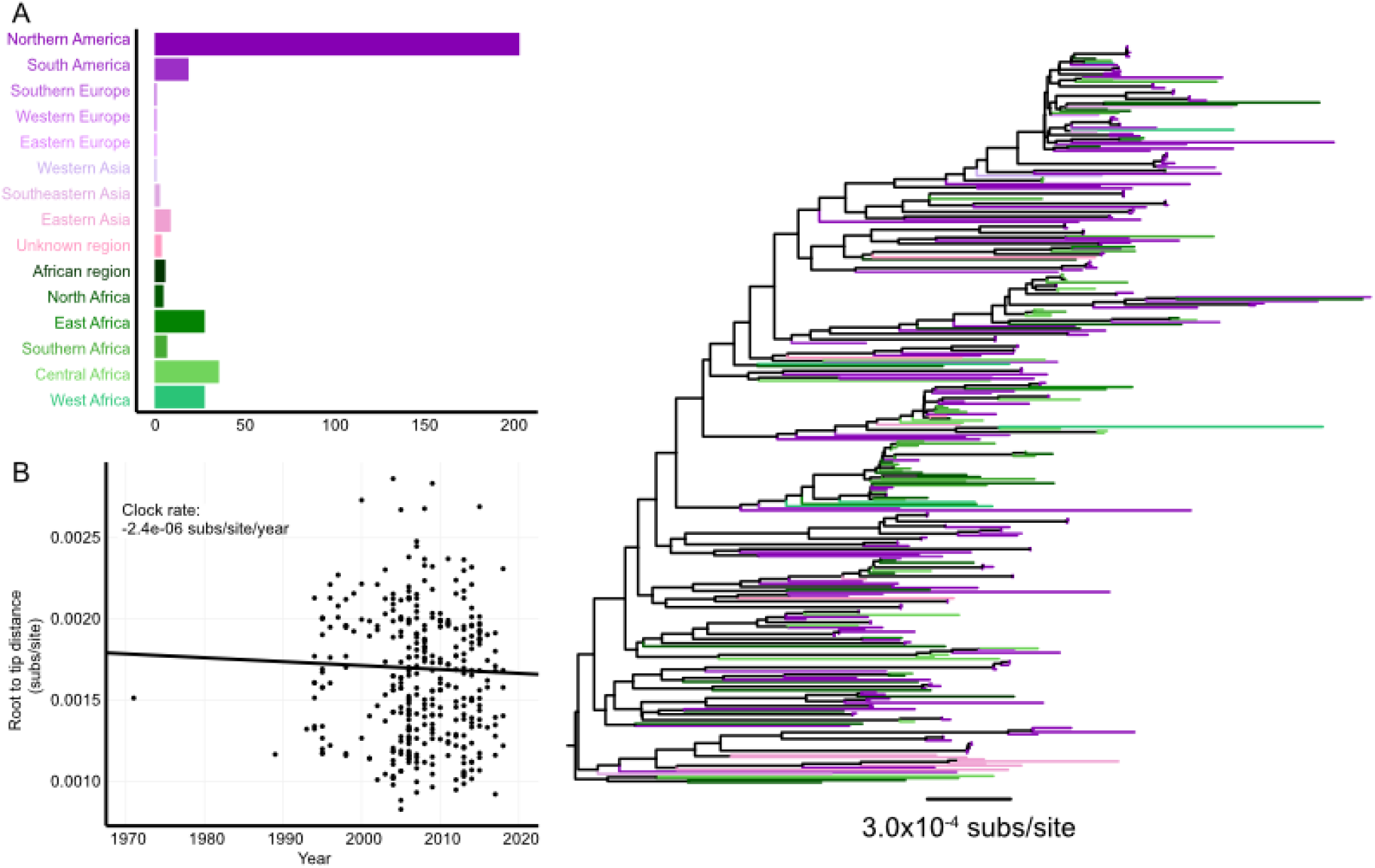
Phylogenetic analysis of subset of HSV-2 worldwide lineage. (A) HSV-2 dataset of 353 sequences in out of Africa clade of worldwide lineage, by number of samples from each region. (B) ML phylogenetic tree inferred with GTR+F+R4 model of sequences described in panel A. Color indicates region of sampling for tip. (C) Root to tip distance of ML tree verse sampling year, with regression estimate of clock rate of -2.4×10-6 substitutions/site/year.

**Fig. S2.**
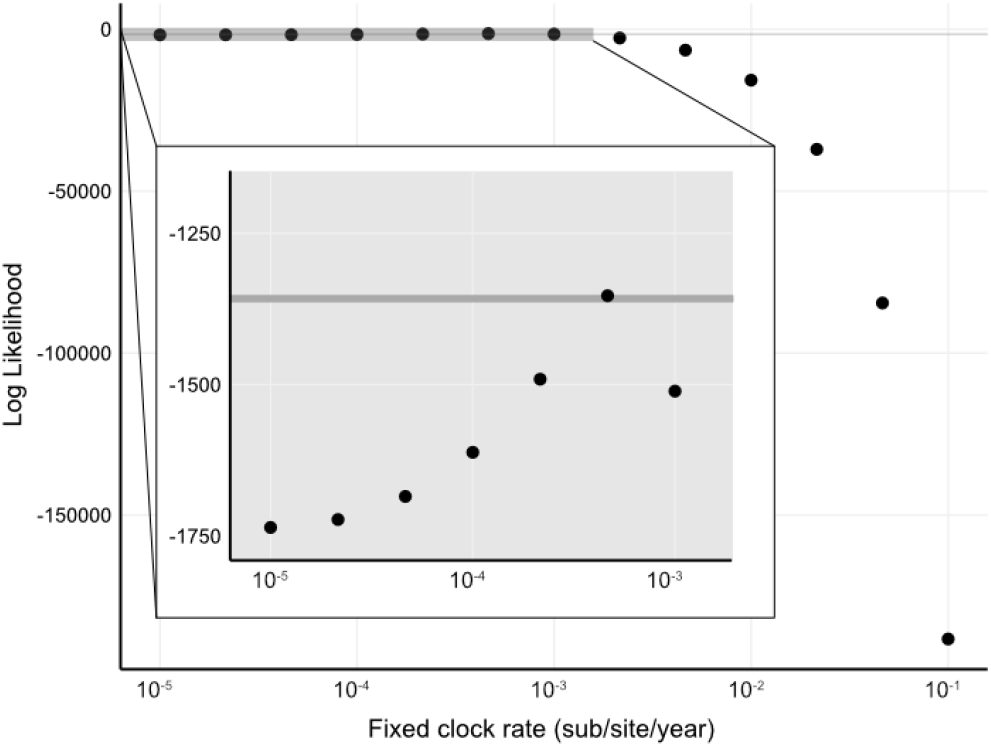
Shape of likelihood surface for clock rate of ebolavirus. Log likelihood of ebolavirus time tree at fixed clock rates estimated using input ML tree (1) of 1610 Ebola genomes, black line is maximum estimated likelihood, grey shaded box is the area shown in insert. (Insert) Black line is maximum estimated likelihood at rate 4.6×10-4 substitutions/site/year, dark grey box is range from maximum estimated likelihood to 10 points below.

**Fig. S3.**
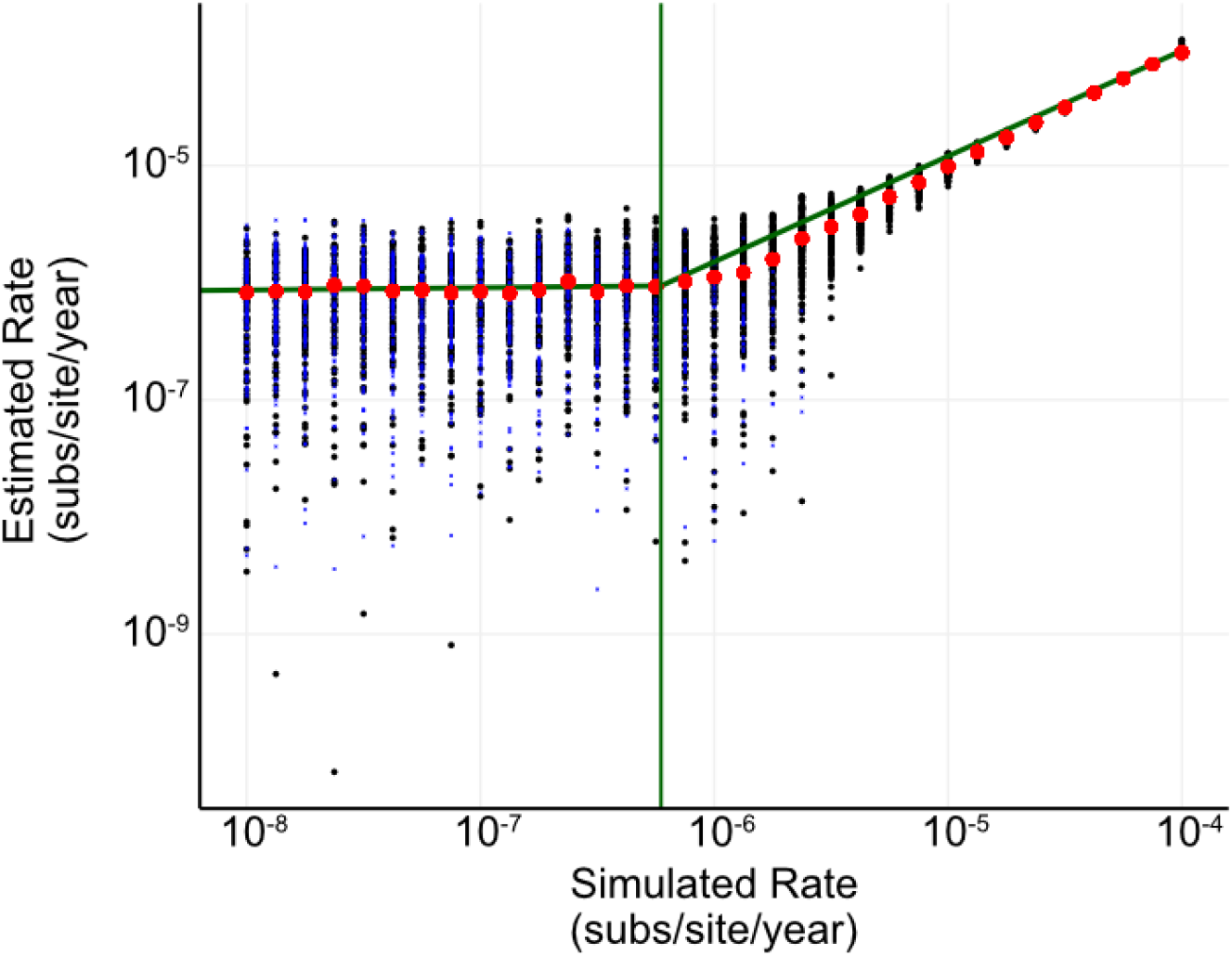
Breakpoint analysis of clock rate inference on simulated data. Estimates of rate for a fixed input tree with simulated sequences and empirical dates. Black dot represents estimates of positive rates, while blue ‘x’ represent the absolute value of estimates of negative rates, and the mean of absolute value for each fixed rate is indicated by red dot. Sloped green segments represent break point regression of absolute value of estimated rates. Vertical green line represents break point at 5.6×10-7 substitutions/site/year (R segmented). 200 replicates were generated under GTR+Γ4 model with fixed rates 1×10-8 to 1×10-4 substitutions/site/year.

**Fig. S4.**
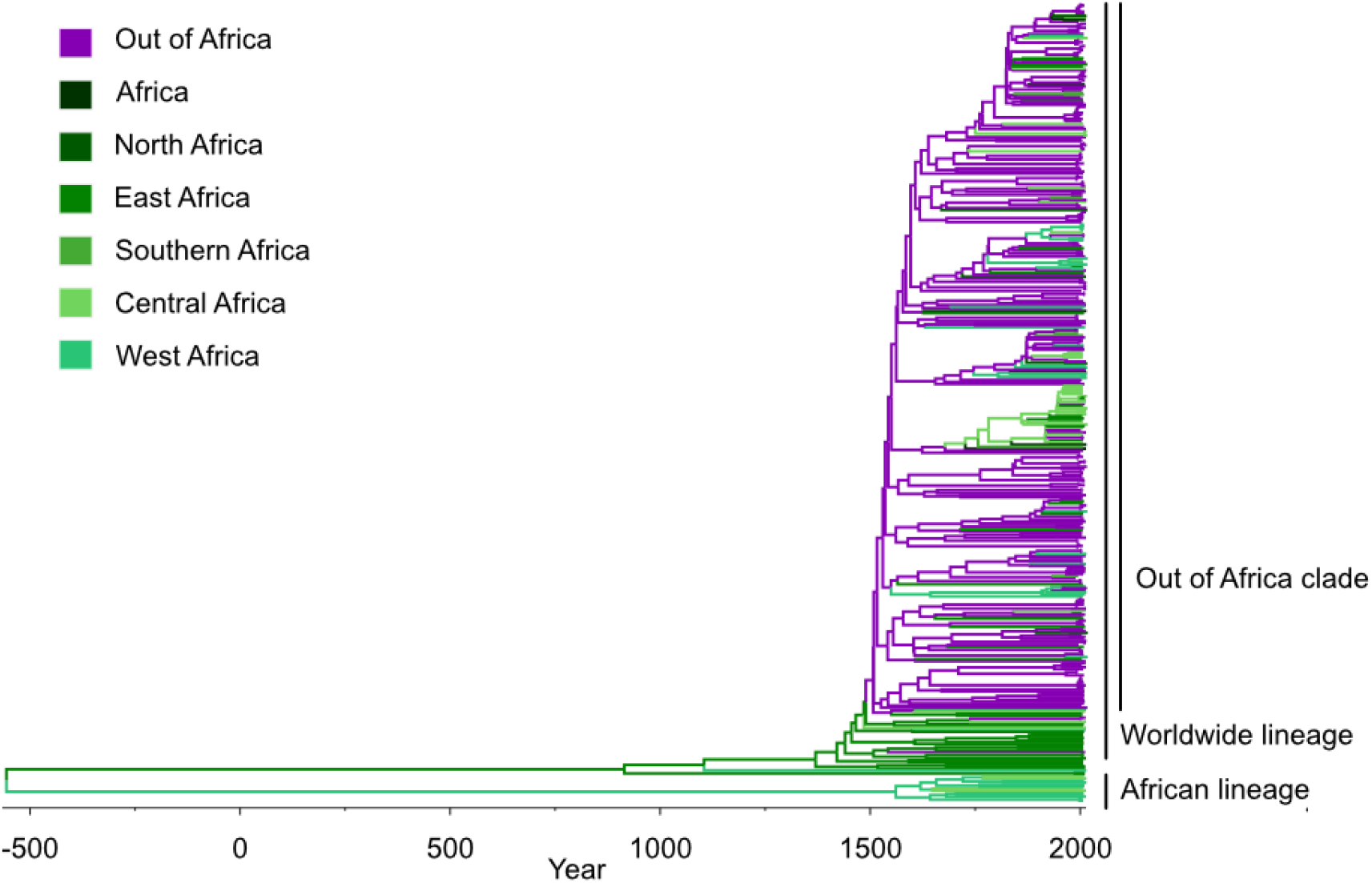
Phylogeography of HSV-2 supports East Africa as origin of out-of-Africa dispersal. Time tree inferred with the fastest rate supported by any clock analysis, 2×10-6 substitutions/site/year. ML tree topology scaled to time tree, colored by plurality of support in mugration analysis for each region. Time scale represents the calendar year; negative years are BCE.

**Fig. S5.**
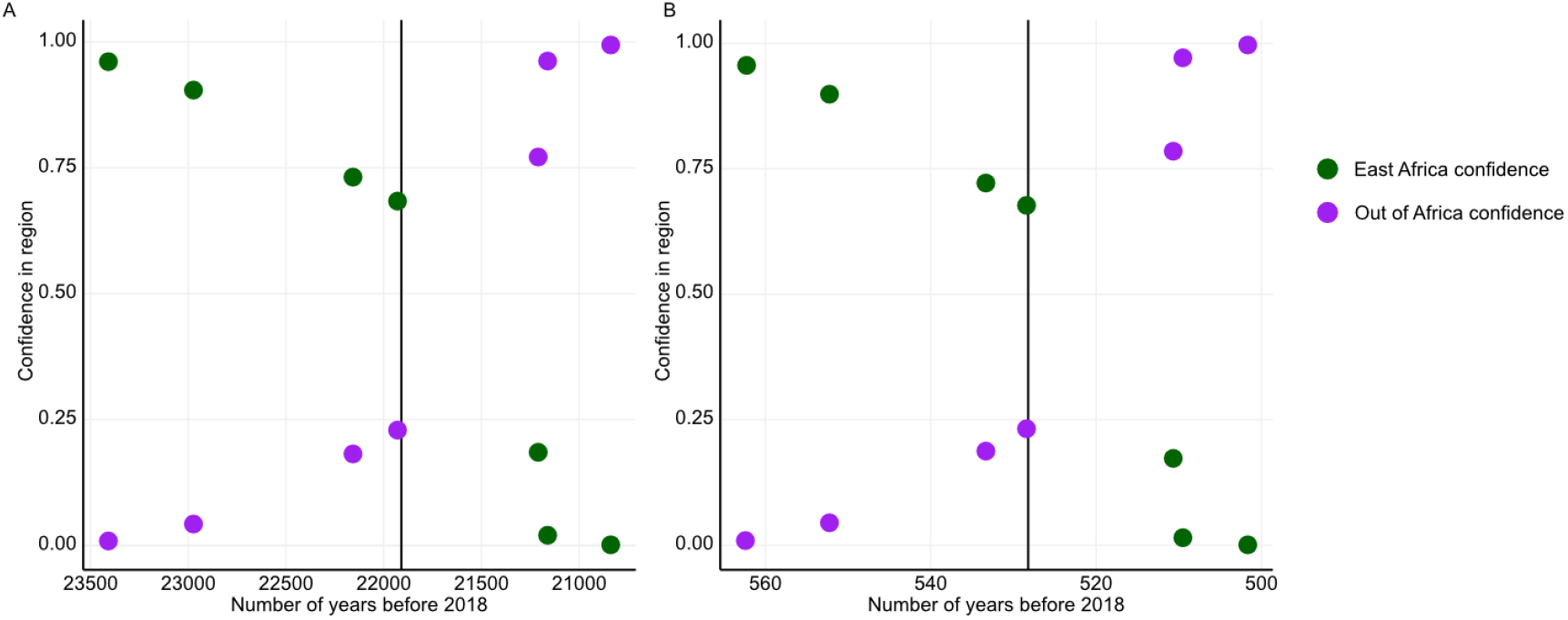
Internal nodes in backbone of tree around out of Africa migration (within the worldwide lineage). Confidence in East Africa placement (green) and out of Africa placement (purple), by number of years before 2018, for time trees with rate 5.8×10-8 (A) and 2.4×10-6 (B) substitutions/site/year. Vertical bar indicates average age of nodes with ages weighted by 1-absolute value(East Africa confidence - out of Africa confidence).

**Table S1.** Age of root and out-of-Africa dispersal across clock rates for robustness analysis of geographic partitioning.

| Partitioning scheme | Substitution rate | Root age<sup>1</sup> | Maximum support for East Africa migration | Rate support in simulations<sup>2</sup> |
| --- | --- | --- | --- | --- |
| All subregions | 1.0E-08 | 122447 | 0.976 | Not rejected |
| All subregions | 1.3E-08 | 91470 | 0.977 | Not rejected |
| All subregions | 1.8E-08 | 68746 | 0.976 | Not rejected |
| All subregions | 2.4E-08 | 51445 | 0.977 | Not rejected |
| All subregions | 3.2E-08 | 38414 | 0.977 | Not rejected |
| All subregions | 4.2E-08 | 28910 | 0.978 | Not rejected |
| All subregions | 5.6E-08 | 21641 | 0.977 | Not rejected |
| All subregions | 7.5E-08 | 16279 | 0.977 | Not rejected |
| All subregions | 1.0E-07 | 12163 | 0.977 | Not rejected |
| All subregions | 1.3E-07 | 9135 | 0.977 | Not rejected |
| All subregions | 1.8E-07 | 6838 | 0.978 | Not rejected |
| All subregions | 2.4E-07 | 5130 | 0.978 | Not rejected |
| All subregions | 3.2E-07 | 3851 | 0.978 | Not rejected |
| All subregions | 4.2E-07 | 2891 | 0.978 | Intermediate |
| All subregions | 5.6E-07 | 2181 | 0.977 | Intermediate |
| All subregions | 7.5E-07 | 1630 | 0.979 | Intermediate |
| All subregions | 1.0E-06 | 1226 | 0.979 | Intermediate |
| All subregions | 1.3E-06 | 923 | 0.981 | Intermediate |
| All subregions | 1.8E-06 | 693 | 0.982 | Intermediate |
| All subregions | 2.4E-06 | 521 | 0.983 | Intermediate |
| All subregions | 3.2E-06 | 396 | 0.984 | Rejected by all |
| All subregions | 4.2E-06 | 299 | 0.986 | Rejected by all |
| All subregions | 5.6E-06 | 226 | 0.988 | Rejected by all |
| All subregions | 7.5E-06 | 173 | 0.989 | Rejected by all |
| All subregions | 1.0E-05 | 132 | 0.992 | Rejected by all |
| All subregions | 1.3E-05 | 102 | 0.994 | Rejected by all |
| All subregions | 1.8E-05 | 80 | 0.995 | Rejected by all |
| All subregions | 2.4E-05 | 64 | 0.997 | Rejected by all |
| All subregions | 3.2E-05 | 54 | 0.997 | Rejected by all |
| All subregions | 4.2E-05 | 52 | 0.993 | Rejected by all |
| All subregions | 5.6E-05 | 50 | 0.967 | Rejected by all |
| All subregions | 7.5E-05 | 49 | 0.824 | Rejected by all |
| All subregions | 1.0E-04 | 49 | 0.437 | Rejected by all |
| Out of Africa and African merged | 1.0E-08 | 124370 | 0.985 | Not rejected |
| Out of Africa and African merged | 1.3E-08 | 92838 | 0.986 | Not rejected |
| Out of Africa and African merged | 1.8E-08 | 69776 | 0.985 | Not rejected |
| Out of Africa and African merged | 2.4E-08 | 52233 | 0.986 | Not rejected |
| Out of Africa and African merged | 3.2E-08 | 38987 | 0.985 | Not rejected |
| Out of Africa and African merged | 4.2E-08 | 29342 | 0.986 | Not rejected |
| Out of Africa and African merged | 5.6E-08 | 21961 | 0.986 | Not rejected |
| Out of Africa and African merged | 7.5E-08 | 16528 | 0.985 | Not rejected |
| Out of Africa and African merged | 1.0E-07 | 12350 | 0.985 | Not rejected |
| Out of Africa and African merged | 1.3E-07 | 9271 | 0.985 | Not rejected |
| Out of Africa and African merged | 1.8E-07 | 6943 | 0.986 | Not rejected |
| Out of Africa and African merged | 2.4E-07 | 5206 | 0.986 | Not rejected |
| Out of Africa and African merged | 3.2E-07 | 3911 | 0.986 | Not rejected |
| Out of Africa and African merged | 4.2E-07 | 2937 | 0.986 | Intermediate |
| Out of Africa and African merged | 5.6E-07 | 2214 | 0.985 | Intermediate |
| Out of Africa and African merged | 7.5E-07 | 1655 | 0.985 | Intermediate |
| Out of Africa and African merged | 1.0E-06 | 1245 | 0.985 | Intermediate |
| Out of Africa and African merged | 1.3E-06 | 937 | 0.985 | Intermediate |
| Out of Africa and African merged | 1.8E-06 | 704 | 0.985 | Intermediate |
| Out of Africa and African merged | 2.4E-06 | 530 | 0.984 | Intermediate |
| Out of Africa and African merged | 3.2E-06 | 403 | 0.984 | Rejected by all |
| Out of Africa and African merged | 4.2E-06 | 304 | 0.984 | Rejected by all |
| Out of Africa and African merged | 5.6E-06 | 230 | 0.984 | Rejected by all |
| Out of Africa and African merged | 7.5E-06 | 176 | 0.983 | Rejected by all |
| Out of Africa and African merged | 1.0E-05 | 134 | 0.984 | Rejected by all |
| Out of Africa and African merged | 1.3E-05 | 104 | 0.984 | Rejected by all |
| Out of Africa and African merged | 1.8E-05 | 81 | 0.986 | Rejected by all |
| Out of Africa and African merged | 2.4E-05 | 65 | 0.988 | Rejected by all |
| Out of Africa and African merged | 3.2E-05 | 55 | 0.989 | Rejected by all |
| Out of Africa and African merged | 4.2E-05 | 52 | 0.977 | Rejected by all |
| Out of Africa and African merged | 5.6E-05 | 50 | 0.939 | Rejected by all |
| Out of Africa and African merged | 7.5E-05 | 49 | 0.827 | Rejected by all |
| Out of Africa and African merged | 1.0E-04 | 49 | 0.551 | Rejected by all |

<sup>1</sup>Years prior to most recently sampled virus, 2018
<sup>2</sup> Support described in Fig. 5

## Notes

### Competing Interest Statement

The authors have declared no competing interest.

